# Transcriptomic Dynamics of a non-coding trinucleotide repeat expansion disorder SCA12 in iPSC derived neuronal cells: signatures of interferon induced response

**DOI:** 10.1101/201137

**Authors:** Deepak Kumar, Parashar Dhapola, Ashaq Hussain, Rintu Kutum, Achal K. Srivastava, Mitali Mukerji, Odity Mukherjee, Mohammed Faruq

## Abstract

Spinocerebellar ataxia type-12 (SCA12) is a neurological disorder that exhibits a unique progressive tremor/ataxia syndrome induced by triplet (CAG) repeat expansion in 5’ UTR of *PPP2R2B*. SCA12 is one of the most prominent SCA-subtype in India and till date no appropriate disease models have been described. Our aim was to establish human iPSC derived neuronal cell lines of SCA12 and study transcriptomic level alterations induced by CAG expansion. For translational application, peripheral blood transcriptomics of SCA12 patients was also performed. Lymphoblastoid cell lines of three SCA12 patients were reprogrammed to iPSCs and then re-differentiated into pan-neuronal lineage. RNA-sequencing based comparative transcriptomics was performed for disease and control cell lineages. Microarray based transcriptomic profiling of peripheral blood of SCA12 patients was performed in a case/control (n=15/9) design. We have successfully created human neuronal cell lines of SCA12 patient as exhibited by their molecular profiling. Differential expression analysis of RNA-Seq data has shown enrichment for type-I interferon signaling and other relevant cellular processes in SCA12-neurons. At the splice-isoform level, we observed an upregulation of expanded CAG containing non-coding transcript of *PPP2R2B*. Peripheral blood transcriptomics analysis and targeted validation of RNA-Seq data has allowed us to identify inflammatory signatures as potential markers of molecular pathology in SCA12. Our study has allowed us to establish first iPSC based neuronal cell lines of SCA12. We have identified pro-inflammatory signatures in SCA12-neurons suggestive of a dsRNA mediated activation of interferon signaling and that corroborates with the emerging evidence of neuronal atrophy due to neuro-inflammation in common neurodegenerative diseases. This study involved development of an iPSCs derived neuronal cells of SCA12 and look through signatures of neurodegeneration by whole RNA sequencing. This model sheds light upon key role of RNA mediated induced response in Interferon signaling for neurodegeneration.

## Introduction

SCA12 is an autosomal dominant trinucleotide repeat expansion disorder clinically characterized by cerebro-cerebellar degeneration (i.e. progressive action tremor of hands, gait ataxia, dysarthria, head tremor etc.) and variable onset of other pyramidal and extrapyramidal features during the course of disease progression (Holmes et al., 1999). The underlying genetic defect in SCA12 is the expansion of CAG repeats (≥43) in 5’ region of *PPP2R2B* (Srivastava et al., 2017). This gene encodes a regulatory subunit of the heterotrimeric enzyme serine/threonine protein phosphatase 2A (PP2A) which is pivotal to essential cellular functions and role in cancer, neurodegeneration and inflammatory diseases (Janssens and Goris, 2001; Sangodkar et al., 2016). *PPP2R2B* transcribes into several repeat containing and non-repeat (non-TNR) containing splice variants that differ at 5’ sequence regions and encode protein isoforms with variable N-terminal, thus affecting the localization and substrate specificity of PP2A enzyme (Strack et al., 1998; Dagda et al., 2003). In earlier reports the role of non -TNR isoform of *PPP2R2B* has shown to induce apoptotic cell death in cellular models and mitochondrial fragmentation with cristae disruption in fly model of SCA12 (Cheng et al., 2009; Wang et al., 2011). But the molecular mechanism by which CAG repeat expansion in PPP2R2B induces neurodegeneration is largely unknown. Clinical manifestations of SCA12 include features that are unique to this disease (e.g. action tremor in hands) as well as those overlapping with other cerebello-degenerative disorders (such as gait ataxia and pyramidal features). At the phenotypic level SCA12 is closely connected with other neurodegenerative disorders like Spinocerebellar ataxia type 1-3 & 17, Huntington’s disease and Parkinson’s disease. Fragile X-associated Tremor/Ataxia Syndrome (FXTAS), which is also considered a phenocopy of SCA12, is caused by TNR expansion of CGG repeats in the 5’UTR of *FMR1,* resulting into RNA toxicity (Hagerman, 2013; Faruq et al., 2014). RNA mediated toxicity has been implicated in many TNR diseases caused by repeat expansion either in the coding or non-coding region (Krzyzosiak et al., 2012). A similar scenario can be anticipated in SCA12 where the CAG repeat expansion in a non-coding region of *PPP2R2B.* Till date, an appropriate disease model to study the mechanism of CAG expansion mediated SCA12 pathogenesis is lacking. Human-induced-pluripotent-stem cell (HiPSC) offers to be a promising model for elucidating the pathology of several human disorders and their usage gaining momentum owing to its ability to model the disease in patients own genetic background (Sterneckert et al., 2014). To study the mechanistic role of CAG repeat expansion in SCA12 pathogenesis, here we describe for the first time, the generation of iPSCs from SCA12 patients, followed by the cellular reprogramming of these SCA12-iPSCs into neurons. In order to identify the transcriptional changes induced by the mutation in the disease condition, we have performed whole transcriptome analysis by RNA-sequencing in patient iPSC-derived neuronal and other cell lineages and then systematically analyzed the changes in *PPP2R2B* transcripts level. We have corroborated these observations with insights from the whole blood transcriptomic profile of SCA12 patients so as to understand the aberrant transcriptional milieu in diseased individuals. Our results show enrichment of neurodegeneration pathways in the RNA-Seq data and aberrant expression of *PPP2R2B*. Surprisingly we find an altered expression of genes related to the interferon signaling pathway in both iPSC-derived neurons and peripheral blood of SCA12 patients.

## Materials and methods

### Subjects and study approval

Patients were clinically and genetically confirmed for the presence of SCA12 at All India Institute of Medical Sciences (AIIMS) before their recruitment into this study. For iPSC generation, blood samples of three unrelated patients and control were drawn. For genome wide expression study using microarray, peripheral blood was collected from 15 unrelated patients [Male/ Female; 12/3, mean age= 53.6 ± 9.1 years, mean expanded CAG length =57.8 ± 6.1] and nine controls [Male/ Female; 2/7, mean age= 51.7 ± 5.7 years]. Additionally seven other SCA12 patients [Male/Female; 9/3, mean age= 54.5 ± 9.8 years, mean expanded CAG length = 56.3 ± 6.3] and five controls were included in this study for the validation of RNA-Seq data Supplementary file 1: Table 1. The study was approved by the Institute Ethics Committee & Institutional Committee for Stem Cell Research, AIIMS, New Delhi (Ref no. IEC/NP-26/2013 & IC-SCRT/19/14R) and the Human Ethics committee of CSIR-IGIB for GENCODE (BSC0123). Informed written consent was obtained from each subject. The samples were collected in strict compliance with the ethical guidelines of Indian Council of Medical Research (ICMR).

**Table 1:** **Enriched biological process in SCA12 patient derived neurons, predicted by Database for Annotation, Visualization and Integrated Discovery (DAVID).**

| ID | Term | Count | P-Value | FDR |
| --- | --- | --- | --- | --- |
| GO:0060337 | Type I interferon signaling pathway | 16 | 2.6 E-9 | 4.6E-6 |
| GO:0007155 | Cell adhesion | 40 | 1.5E-7 | 8.8E-5 |
| GO:0030198 | Extracellular matrix organization | 23 | 9.0E-7 | 1.6E-3 |
| GO:0006504 | Basement membrane | 14 | 2.0E-6 | 2.7E-3 |
| GO:0001525 | Angiogenesis | 24 | 2.3E-6 | 4.0E-3 |
| GO:0045071 | Negative regulation of viral genome replication | 10 | 6.0E-6 | 1.1E-2 |
| GO:0045944 | Positive regulation of transcription from RNA polymerase II promote | 58 | 4.9E-5 | 2.1E-2 |
| GO:0007417 | Antigen processing and presentation of exogenous peptide antigen | 5 | 1.4E-4 | 4.6E-2 |

**GO=Gene ontology; p<0.05; FDR: false discovery rate <0.05**

### Peripheral blood mononuclear cells (PBMCs) isolation, LCL generation and characterization

PBMCs were isolated from three unrelated symptomatic SCA12 patients and one control using ficoll histopaque (Sigma-Aldrich) gradient following manufacturer’s instructions. PBMCs were then transformed to lymphoblastoid cell lines (LCLs) using Epstein Barr virus (EBV) following a published protocol (Frisan et al., 2001). LCLs were maintained in RPMI 1640 medium (Life Technologies) supplemented with 20% FBS (Gibco), 2 mM glutamine and 1X Penicillin & Streptomycin (Life Technologies). Direct immunofluorescence was measured to confirm the presence of B cell subpopulation using CD19 cell surface marker. CD3 staining was also performed to confirm the exclusion of T cells in culture. Cells were washed once with 1 X DPBS, suspended in FACS buffer containing diluted FcR blocking and incubated on ice 10 minutes. Blocking was removed and cells were resuspended in FACS buffer and antibodies labelled with fluorophore (CD19-PerCP-Cy5.5 and CD3-FITC (BD Biosciences) as per manufactures instructions. Cells were then incubated on ice for 30 minutes, washed once with FACS buffer, resuspended in PBS and acquired on FACS Aria (BD Biosciences).

### Induced pluripotent stem cells (iPSCs) generation

The generated LCLs of the three patients and one control were reprogrammed via episomal plasmids harboring reprogramming factors as described previously (Okita et al., 2011). Briefly 0.2 million LCLs were electroporated with 1μg of three reprogramming plasmids viz., pCXLE-hOCT3/4-shP53-F (Addgene), pCXLE-hSK (Addgene) and pCXLE-hUL (Addgene) using Neon device (Invitrogen). Post electroporation, cells were transferred onto gamma irradiated mouse embryonic fibroblast feeder layer and cultured in iPSC medium containing KnockOut DMEM, 20% KnockOut Serum Replacement, 55 mM beta-mercaptoethanol, 10 mM nonessential amino acids, 2 mM L-glutamine, 1X Penicillin & Streptomycin, 10 ng/ml recombinant human bFGF (PeproTech) supplemented with 50 μg/ml L-ascorbic acid (Sigma-Aldrich), and 0.5 mM sodium butyrate (Sigma-Aldrich). Reagents were obtained from Life Technologies except stated otherwise. As early as 12–15 days after electroporation, typical human embryonic stem (ES) cells like colonies were observed. Once the colonies grew, they were mechanically passaged onto inactivated mouse embryonic fibroblasts (MEF) feeder layer and propagated in iPSC medium lacking sodium butyrate and ascorbic acid. Karyotyping was carried out using G banding to check any gross chromosomal abnormalities. Established iPSC lines were characterized with respect to their pluripotent properties. In-vitro pluripotency assessment was done through embryoid body (EB) formation. For this iPSC colonies were disintegrated (clumps containing 200-300 cell) and transferred into low adherent dishes (Corning) and cultured in EB medium (iPSC medium without basic FGF) till day 4. Then total RNA was isolated from these EBs using RNeasy Mini Kit (Qiagen) and RNA was converted into cDNA using High-Capacity cDNA Reverse Transcription Kit (Applied Biosystems). Reverse transcriptase PCR was performed for different germ layer markers (ectoderm, endoderm and mesoderm) (Supplementary file 1: Table 2). The amplified products were checked on 2% agarose gel using UV trans-illuminator documentation system (Bio Rad).

### Neural stem cells (NSCs) generation and neuronal differentiation

Of all the generated cell lines, one of the patient iPSC (IG0002iPSSCA12) and the control iPSC (NC0001iPSCCE) were differentiated towards pan-neuronal lineage using a published protocol (Zhang et al., 2001) with slight modifications. The iPSCs were first converted into embryoid bodies (EBs). After 4-5 days of culture in the neural induction medium (DMEM/F-12 with GlutaMax supplemented with MEM non-essential amino acids (1X), N2 supplement (1X), B27 supplement without vitamin A (2X), basic FGF (20 ng/ml) (PeproTech), EGF (10 ng/ml (PeproTech) Heparin (2μg/ml) (Sigma) and Penicillin-Streptomycin (1X), EBs generated rosette-like structures at their center. The patches of rosettes were marked, manually cut and cultured on laminin-coated dishes in neural expansion medium to obtain a homogeneous population of NSCs. Both patient (IG0002NSCSCA12) and Control (NC0001NSCCE) NSCs were immunostained for neuronal stem cell marker, NESTIN.

NSCs were differentiated into neurons as previously described (Denham and Dottori, 2009; Trujillo et al., 2009) with some modifications. Briefly NSCs were plated at a density of 2-3x10^4^ cells/cm^2^ on Poly-D-lysine/laminin-coated dishes and culture in neural differentiation medium containing neural basal medium supplemented with N2 (1X) and B27 (2X) (with vitamin A) and Penicillin-Streptomycin (1X). On day 7 of differentiation, dibutyryl cAMP (0.5 mM) (Sigma) was added daily for three consecutive days. To obtain mature neurons, cells were grown for 6-7 weeks in the neuronal differentiation medium. Both patient (IG0002NEURONSCA12) and control (NC0001NEURONCE) neurons were characterized for the neuronal markers including TUJ1 (Beta III tubulin) and MAP2 and CAG repeat length of both patient and control’s NSCs and Neurons were identified. Reagents were obtained from Life Technologies except stated otherwise.

### Immunostaining

Cells were fixed with 4% paraformaldehyde (PFA) for 20 minutes followed by permeabilization with 0.1% Triton-X 100 for 10 minutes. The cells were kept in blocking solution (1% BSA) for one hour at room temperature, incubated overnight at 4°C with diluted primary antibodies. The cells were washed thrice with 1X HBSS (5 minutes per wash) and incubated for 1 hour with diluted secondary antibodies at room temperature. Counter staining for nuclei was done with Hoechst 33342 for 20 minutes at room temperature. Slides were mounted in DABCO and imaging was done using epifluorescence microscope (Nikon Eclipse TE2000-E-PFS, Japan) (Supplementary file 1: Table 3).

### RNA isolation, cDNA synthesis and quantitative PCR (qPCR)

Total RNA was isolated from cells using RNeasy Mini Kit (Qiagen). To remove any genomic DNA contamination from isolated RNA samples, DNase treatment was given using TURBO DNA-free Kit (Ambion). Additionally to study the expression pattern of *PPP2R2B* in different brain regions, RNA of different parts of brain (viz. cerebellum, frontal lobe, parietal lobe, medulla oblongata, pons, putamen, substantia nigra) from healthy individuals were procured from CloneTech. cDNA was synthesized from 1μg total RNA using High Capacity cDNA Reverse Transcription Kit (Applied Biosystems). Primers were designed using an online tool Primer3 (http://bioinfo.ut.ee/primer3-0.4.0/primer3/) (Supplementary file 1: Table 2). qPCR was performed in three technical triplicates on Roche 388 platform (Light Cycler 480 II) using SYBR Green Chemistry (KAPA Biosystems). Relative gene expression was calculated using published methods (Livak and Schmittgen, 2001; Schmittgen and Livak, 2008).

### RNA sequencing (RNA-Seq)

RNA-Seq libraries were prepared with 1 μg RNA using TruSeq RNA sample preparation kit. Cluster generation was carried out on cBot using Illumina SBS kit.v3 protocol. Libraries were sequenced on Illumina HiSeq 2500 (Illumina). Generated bcl files were converted into FASTQ using CASAVA v1.8. Trimming and quality filtering was performed using FASTX tool kit (http://hannonlab.cshl.edu/fastx_toolkit/download.html) and FASTQC (http://www.bioinformatics.babraham.ac.uk/projects/download.html#fastqc). The sequenced reads were further mapped to the reference transcriptome (hg38) using Tophat v2.0.5 allowing a maximum of two mismatches. In order to quantify the gene expression in terms of Fragments Per Kilobase of transcript per Million mapped reads (FPKM) was performed using Cufflinks v2.0.2. FPKM were further used by Cuffdiff v2.0.2 to estimate differential expression of genes (DEG) between patients and controls.

### Genome wide expression profiling using microarrays

For genome wide expression analysis using microarray, total RNA was isolated from the peripheral blood of SCA12 patients (n=15) and controls (n=9) using trizol. To reduce the variability, blood collection was done two hours after a light meal for all the subjects and within two hours, RNA extraction procedure was started. For the microarray experiment biotin labelled cRNA samples were generated using TotalPrep RNA amplification kit (Illumina) and hybridized to human WG-6 v3.0 Expression BeadChip array. The Bead Chip was scanned on Illumina iScan System and raw intensity data was extracted without background correction and normalization using GenomeStudio Software Illumina Inc. Probes with detection p-value of 0.05 in at least 3 samples out of 6 in a given array were retained for further analysis. Background correction and quantile normalization was performed using limma package in R. Additionally, we also checked for batch effect using various methods like SNM (supervised normalization method) and ComBat (sva package). To check for the probes (genes) with significant differential expression between control and patients, we have applied t-test/ Wilcoxon test (depending on the distribution). Significant differentially expressed genes between two groups were determined after applying a fold change ≥2.0 or ≤**-** 2 with p value cut off < 0.05.

### CAG repeat analysis for detection of expansion mutation

Genomic DNA was extracted from cells using QIAamp DNA Mini Kit (Qiagen). PCR assay was used to amplify CAG repeats region of *PPP2R2B* using primer sequences (Supplementary file 1: Table 2). Fragment analysis was performed on ABI 3730xl Genetic Analyzer and the length of CAG repeats was calculated using GeneMapper (ABI) software after the estimation of the obtained fragment size.

### Statistical analysis

For RNA-Seq data analysis, differentially expressed genes (DEGs) with log2 fold change of ≥1 or < -1 and p-value of < 0.05, with q-value <0.05 were taken for further analysis. For qPCR, the statistical significance was calculated using Student’s t-test. Non-parametric Mann–Whitney rank sum test was applied in case of non-Gaussian distribution. Statistical analysis was performed using GraphPad Prism 5.0 (GraphPad software).

### Availability of data and materials

RNA-Seq data have been deposited in NCBI with accession number SRP110347.

Genome wide expression analysis data for peripheral blood have been deposited with the accession number (GSE101288).

## Results

### Development of iPSC derived neurons of SCA12

We have generated LCLs from one control (NC0001LCLCE) and three unrelated patients (IG0002LCLSCA12, IG0003LCLSCA12, and IG0004LCLSCA12). Immunophenotyping of these lines showed maximum cell population (87-88%) were positive for B cells markers and negative for T cells in cultured population of cells, indicating their B cell origin. Genotyping showed CAG expansion length remained unchanged in these LCLs. (Fig. 1A-C). LCLs were then reprogrammed into iPSCs and established lines of both control (NC0001iPSCCE) and patients (IG0002iPSCSCA12, IG0003iPSCSCA12, IG0004iPSCSCA12) were characterized with respect to their pluripotent properties. All the lines showed typical Embryonic stem cells (ES) like morphology in co-culture with inactivated MEF feeder layer cells. iPSCs cultured in standard stem cell medium exhibited blue fluorescence, characteristics of pluripotent stem cells in primed state as reported previously (Muthusamy et al., 2014). The iPSCs were immunopositive for pluripotency markers (OCT4, SOX2, SSEA4) (Fig. 2A) and alkaline phosphatase (data not shown). All lines had shown normal karyotype (Fig. 2B) and exhibited in-vitro potential to differentiate into three cellular lineages (ectoderm, mesoderm and endoderm) as evident from their ability to form embryoid body and expressed markers of all the three lineages (Fig. 2C). *PPP2R2B*-CAG length estimation showed no gross genotypic aberrations (Fig. 2D). Out of four derived cell lines, one control iPSC (NC0001iPSCCE) and one patient iPSC (IG0002iPSSCA12) were differentiated towards the pan-neuronal lineage. The iPSCs were first converted into NSCs in neural induction medium and the generated NSCs were confirmed to be nestin positive (Fig. 3A and Fig. 3B). NSCs were subsequently differentiated into neurons and characterized using neuronal markers TUJ-1(Beta III tubulin) and MAP2, (Fig. 3C-F). Both NSCs and neurons exhibited length of the CAG expanded allele was same as it was in PBMCs from which lines were derived (9/16 for control line and CAG 14/60, for patient line) (Fig. 3G and 3H).

**FIGURE 2:**
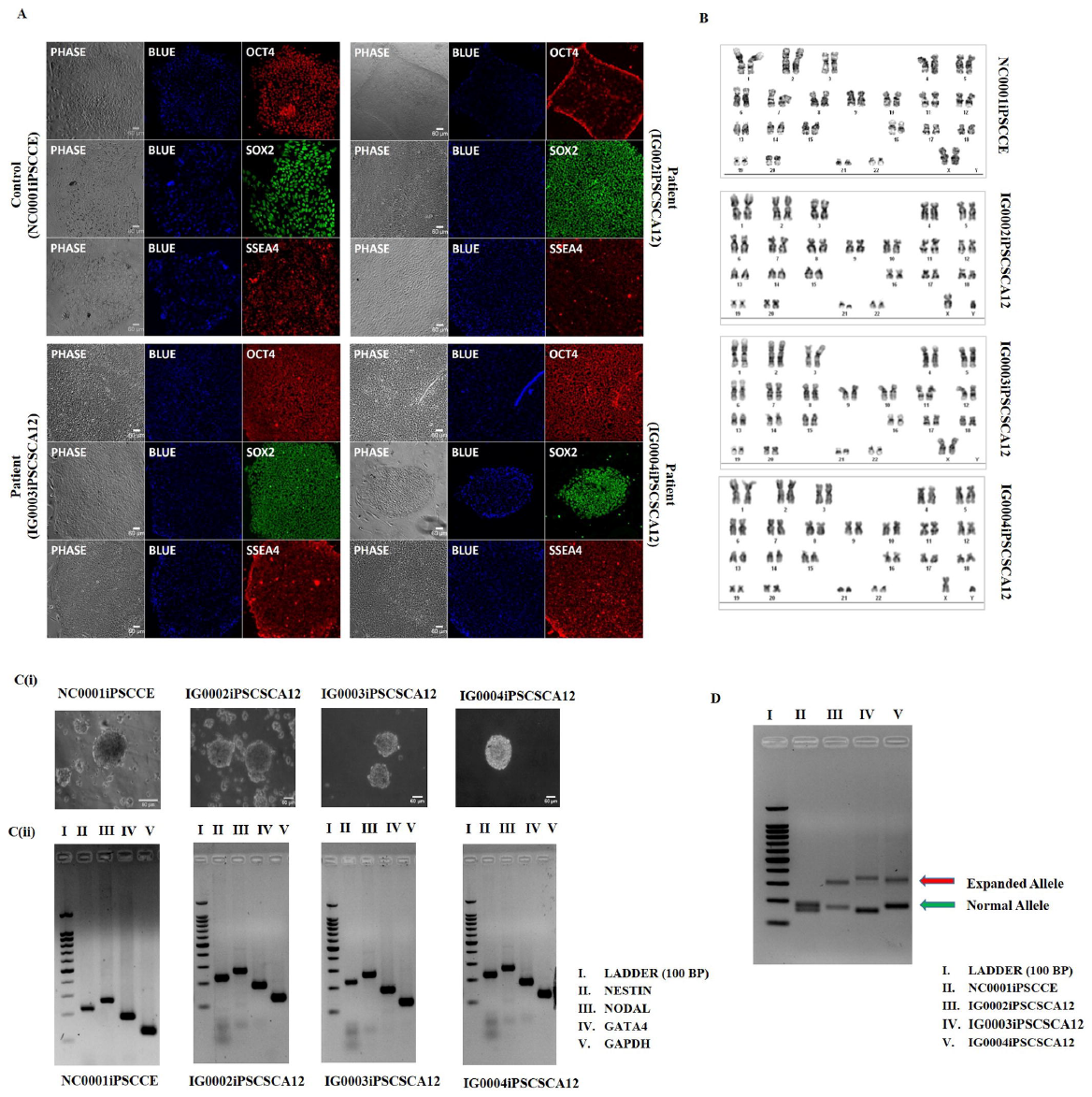
Characterization of induced pluripotent stem cells (iPSCs). **(A, B)** iPSC colonies derived from control (NC0001iPSCCE) and patients (IG0002iPSCSCA12) showing typical embryonic stem cells (ES) like morphology (left panel) and positivity for blue retinyl esters florescence characteristics of pluripotent stem cells in primed state (middle panel). Right panel images indicate immunocytochemistry for pluripotency markers OCT4, SOX2 and SSEA4 (Scale bar, 60 μm). (B) Representative karyotype of all the four iPSC lines derived from control (NC0001iPSCCE) and patients (IG0002iPSCSCA12, IG0003iPSCSCA12 and IG0004iPSCSCA12). (C_(i)) Agarose gel image showing expression of transcripts Nestin (Ectoderm), Nodal (Mesoderm) and GATA4 (Endoderm) in Embryoid body (EBs) generated from SCA 12 patient and healthy control (C-(ii)). (Lane 1= DNA ladder 100 base pair, Lane 2 = Nestin, Lane 3=Nodal, Lane 4= GATA4 and Lane 5= GAPDH, representing as a positive amplification. (D) Agarose gel image showing patients derived iPSC lines are carrying a PPP2R2B CAG heterozygous mutation and no mutation detection in control’s derived iPSCs.

**FIGURE 3:**
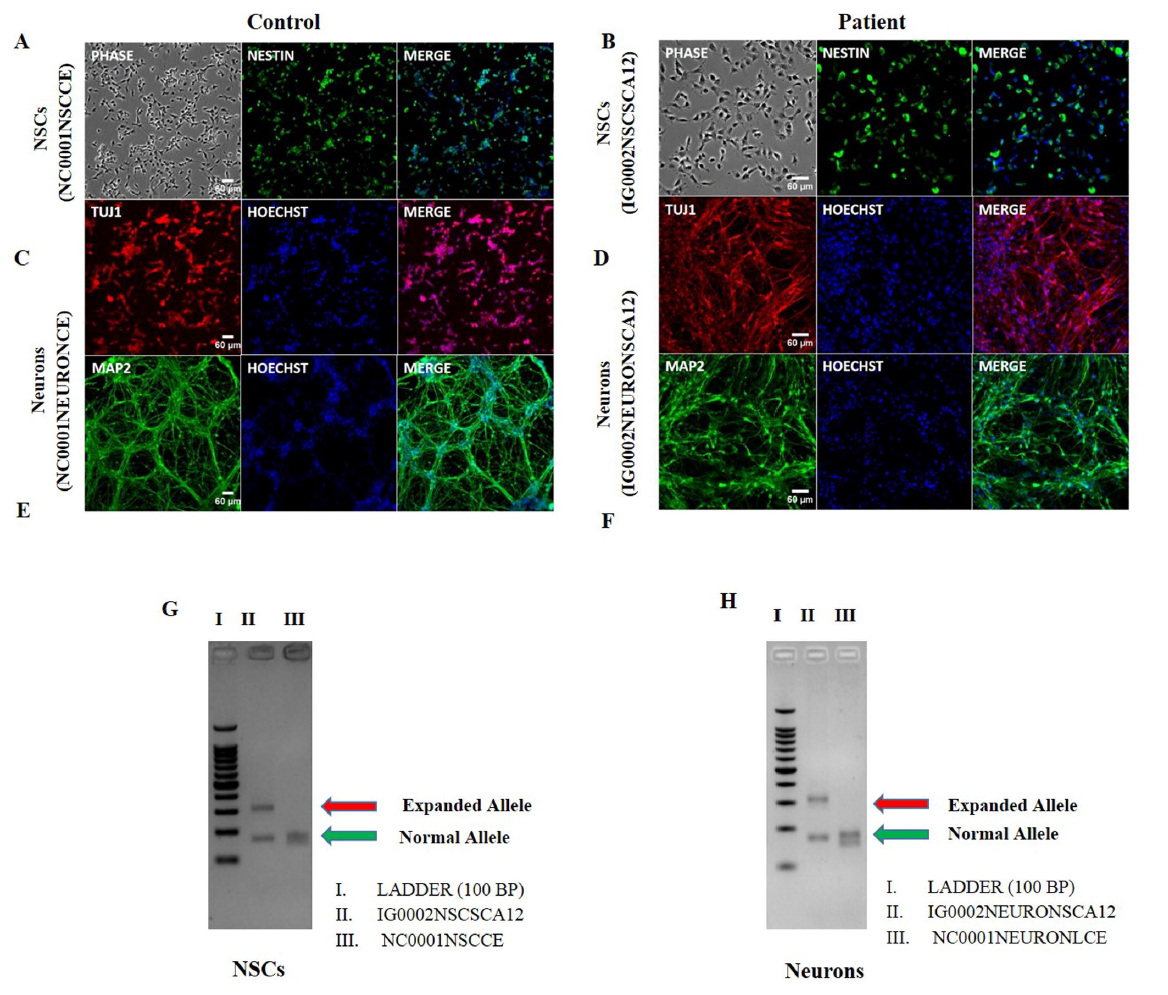
Characterization of neural stem cells (NSCs) and differentiated neurons. (**A, B)** immunocytochemistry for NSC marker NESTIN in control and patient derived lines. Images from **C-F** represent immunocytochemistry for neuronal markers TUJ1 and MAP2 in neurons differentiated from both control and patient derived NSCs. (G, H) Agarose gel image showing patients derived NSCs and neurons carrying a PPP2R2B CAG heterozygous mutation and no mutation detection in control’s derived lines. (Lane 1= DNA ladder 100 base pair) Lane 2 = diseased NSC IG0002NSCSCA12 and diseased neurons IG0002NEURONSCA12. Lane 3= control’s derived NSC (NC0001NSCCE and controls neurons (NC0001NEURONCE).

**FIGURE 1:**
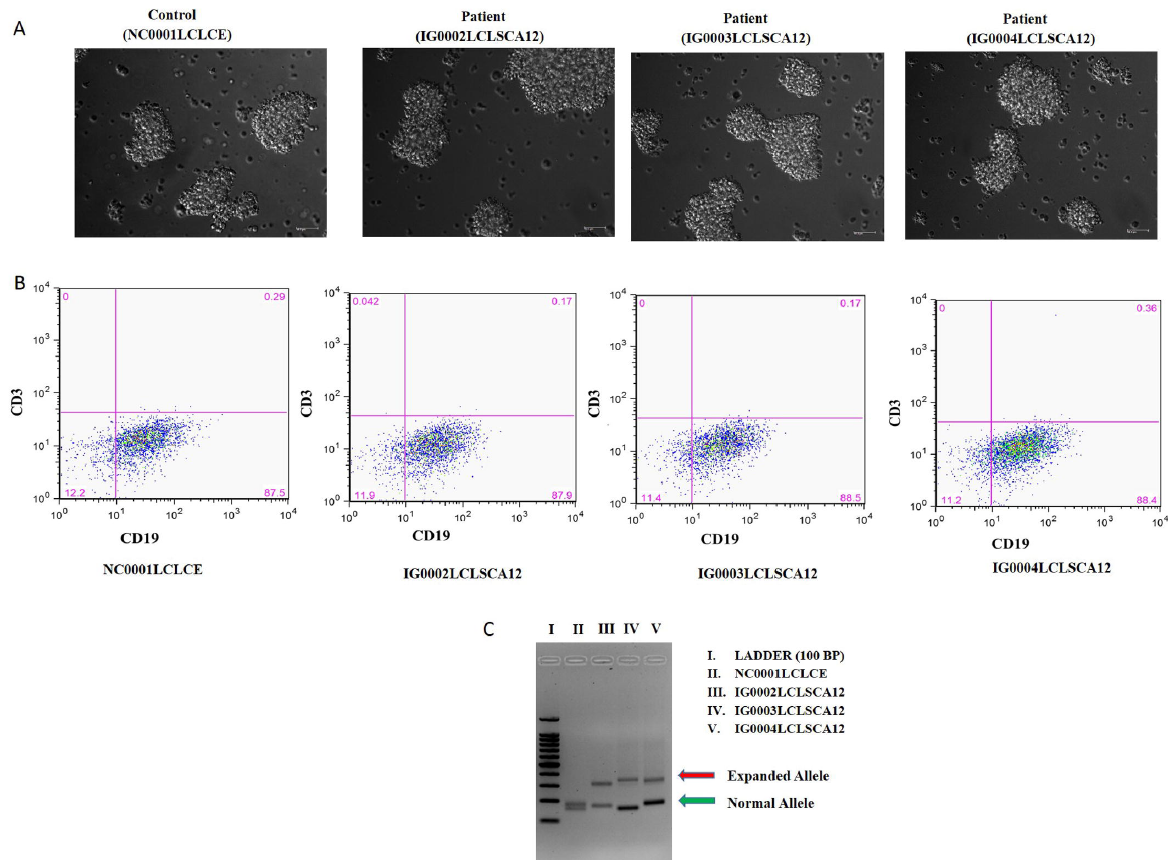
Establishment of lymphoblastoid cell lines (LCLs) NC0001LCLCE, IG0002LCLSCA12, IG0003LCLSCA12, and IG0004LCLSCA14 from PBMCs isolated from control and SCA12 patients’ blood respectively. (A) Morphologies of LCLs showing lymphoblastoid cell aggregates (Scale bar 60 μm) (B) Flow cytometry analysis for B cell marker (CD19) and T cell marker (CD3) (C) Agarose gel image representing SCA12 mutation status in LCLs derived from patients and control.

### Transcriptomic signatures in SCA12 neurons

RNA sequencing (RNA-Seq) was carried out for polyA enriched RNA from cell lines of patients, iPSCs (n=3), neural stem cell (n=1), Neuron (n=1) and for a control of each respective cell line with technical replicates, using TruSeq RNA sample preparation kit (Illumina). Overall we obtained, on an average of 34 Million (M) total processed paired sequence reads per samples, except for one sample (IG0002iPSCSCA12-2) with low sequenced reads. The numbers varied between 12M-82M read pairs. The percentage alignment for sequenced reads was obtained at 90% ± 4 for all the samples (Supplementary file 1: Table 4).

Figure 4A represents the volcano plot representation of transcripts expressed in neurons (volcano plot of iPSC and NSC are not shown). The upregulated/downregulated DEGs (DEG set-I) obtained between disease and control (with cut-off of Log2FC > 1 and < -1 and q value (FDR) ≤ 0.05) for each lineage were following: iPSCs: 556/339, NSCs: 542/828, Neuron: 418/331 genes. We focused upon DEGs from the neuronal lines and further obtained DEG set-II with an additional cut-off of FPKM >1 or cumulative FPKM >10 (for stringency filter). Using this cut off a set of 398 (225 up regulated and 173 down regulated) DEGs were identified from comparison between disease and control neurons. From this DEG set-II, the top 30 up and down regulated genes are represented in Figure 4B. Among these top up/down regulated genes, we observed highest upregulation of *LINC00507*, a long non-coding RNA shown to have brain specific and age dependent expression*. INPP5F, DCN, BACE2 NDFU5S2* and genes linked to inflammation *IFI27, ISG15, HLA-A, HLA-B, IFI16* were among top dysregulated genes. We further explored the role of genes involved in different pathways governing various biological functions. Upregulated genes were found to be involved in interferon signaling, apoptosis, Alzheimer’s disease, Parkinson’s disease, Huntington’s diseases, antioxidant and p53 pathways. On the other hand, downregulated genes were involved in the positive regulation of *ERK1* and *ERK2*, CNS development, NGF signaling and oxidative phosphorylation (Fig. 4C). Validation by qPCR of RNA-Seq data for *IRX1, BACE2, DLK1* and *RBP1* showed concordance in terms of differential expression (Fig. 4D).

**FIGURE 4:**
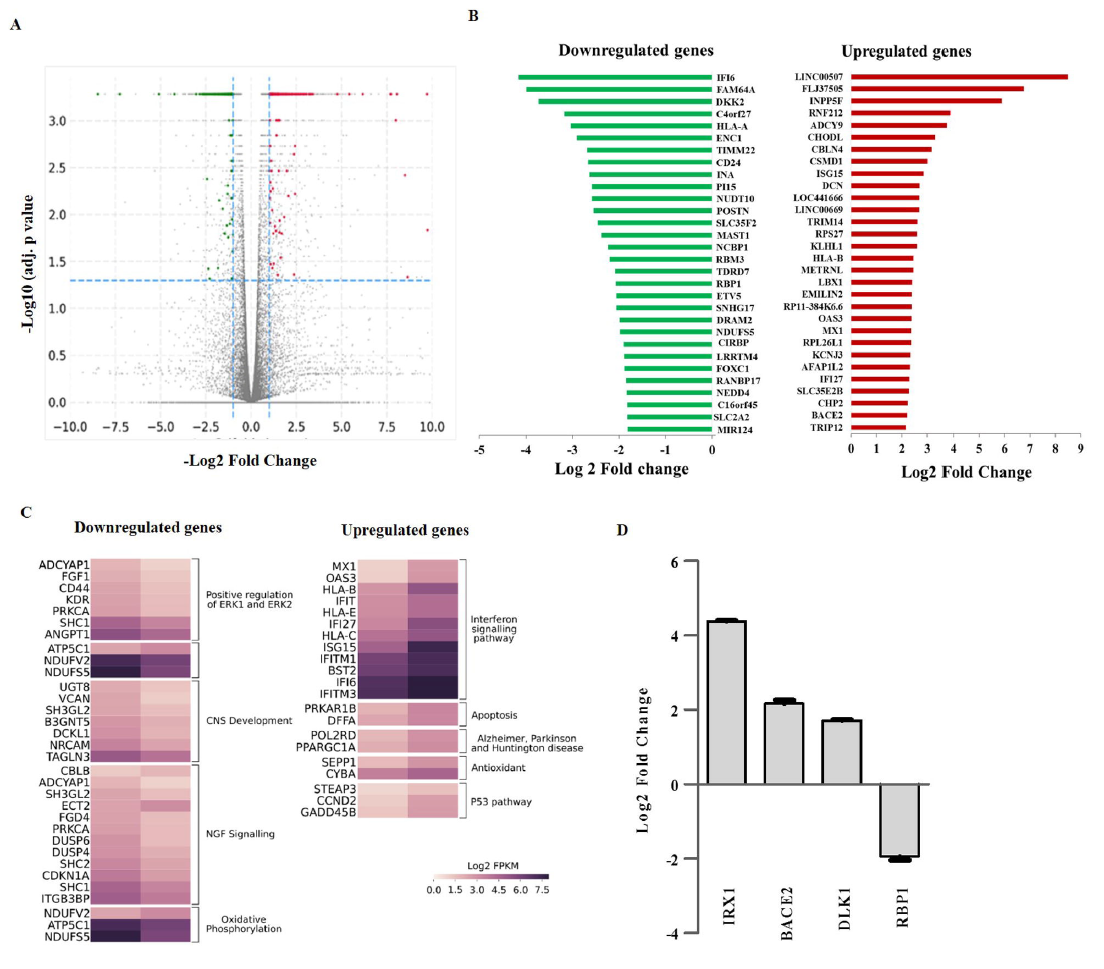
Representation and validation of selected targets through RNA sequencing. **(A)** Volcano plot depicting expressed genes in control and patient derived neurons. For all transcripts log2 fold-change expression values (x axis) are plotted against -log10 corrected p-values (y axis). Horizontal dotted lines represent the thresholds for significance and vertical dotted lines indicate log2 fold change expression. Grey color represents non-significant expression of the transcripts whereas right panel red color denote upregulation and left panel green color represents downregulation of transcripts. **(B)** Representation of top 30 upregulated and downregulated genes differentially expressed in neurons. **(C)** Heat map of differentially expressed genes from top enriched pathways and pathways related to neurodegenerative disorders. **(D)** qPCR based validation of subset of differentially expressed genes between control and patient derived neurons identified by RNA sequencing. GAPDH was used as internal control for normalization. Relative gene expression was calculated using 2 ^-Δ ΔCt^method and results were plotted as log2 fold change in patient derived neurons compared to control.

### Gene ontology (GO) analysis of transcriptome in SCA12 neuron

Further to explore biological function of gene expression data, gene ontology analysis was performed using Database for Annotation, Visualization and Integrated Discovery (DAVID) and Gene Set Enrichment Analysis (GSEA) (Subramanian et al., 2005; Huang et al., 2009) (Supplementary file 2). Functional annotation clustering of all the 749 DEGs using DAVID at FDR of 5% revealed enrichment of 28 gene terms that includes Interferon signaling (14/2, up/down genes), cell adhesion (24/16, up/down genes), ECM matrix (18/5, up/down genes) and calcium ion binding (14/3, up/down genes) terms. Of these enriched 28 term, GO biological term are depicted in Table 1. Additionally, GSEA of our differential expression data have shown top 10 gene sets and top term was response of interferon signaling pathway and immune system. Axon guidance (10/6, up/down) and NGF signaling pathway (6/13, up/down) were identified among other enriched gene set (Supplementary file 2).

In addition we also performed GO analysis for DEGs from RNA-Seq data of iPSC and NSC lineage and observed that interferon signaling was enriched only in the neuronal lineage (Supplementary file 1: Table 5 and 6). To find distribution of differentially expressed genes related to interferon signaling, we performed Venn distribution of interferon enriched genes among these generated iPSCs, NSCs and neurons and found overrepresentation in neurons (Fig. 5A and Supplementary file 1: Table 7). In neurons, among induced interferon response, 14 up regulated genes were *ISG15, HLA-B, OAS1, OAS3, MX1, IFI27, IFI6, SAMHD1, IFIT1, IFITM1, IFITM3, HLA-C, HLA-E* and *BST2* while, *EGR1*, *HLA-A* were downregulated (Fig. 5B). We further validated *OAS1* and *ISG15* in neurons using qPCR and found log 2.8 fold upregulation in expression of *ISG15* and log 2.2 fold for *OAS1* (Fig. 5C).

**FIGURE 5:**
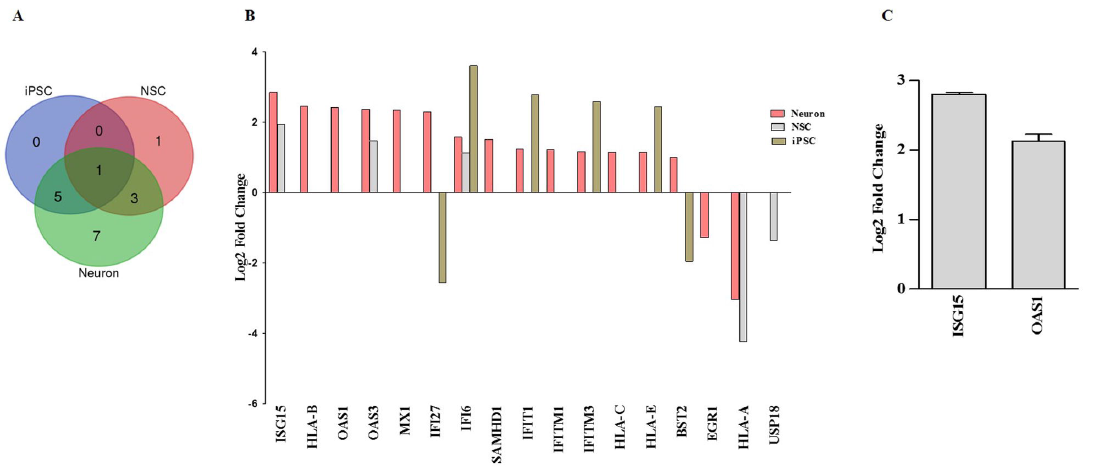
Enrichment of genes related to interferon type 1 signaling pathway in patient derived neurons. **(A)** Venn diagram representing overlap of differentially expressed genes related to interferon type 1 signaling pathway among patient derived iPSC, NSC and neurons compared to their respective control. **(B)** Log2 fold expression of interferon type 1 signaling pathway related genes in iPSC, NSC and neurons identified from RNA sequencing analysis. **(C)** Validation of *OAS1* and *ISG15* expression in patient derived neurons relative to control by qPCR. GAPDH was used as internal control. Relative gene expression was calculated using 2 ^-Δ ΔCt^method and results were plotted as log2 fold change in patient derived neurons compared to control.

### Transcriptomic alteration of genes involved in IFN response and Neurodegeneration

Poly (ADP-ribose) polymerase (PARPs) is a group of stress response family of proteins that have role in diverse functions including DNA damage response and programmed cell death. We found differential expression of *PARP9* and *DTXL3*. PARP9-DTX3L ubiquitin ligase interact with STAT1and targets host histone to promote interferon-stimulated gene expression which induces degradation of pathogen protease (Zhang et al., 2015). We further observed upregulation of genes involved in P53 pathway *STEAP3, THBS1, CCND2. GADD45B PERP* and *CDKN1A*. In our study we also observed upregulation of *DDX58* and *IFIH1* in disease condition. There was also an overlap of altered genes with genes related to neurodegenerative diseases, including Alzheimer’s disease (AD) (hsa05010), Parkinson’s disease (PD) (hsa05012), (hsa05014), Huntington’s disease (HD) (hsa05016) and Prion disease (hsa05020) (Supplementary file 1: Table. 8).

### Similarities of SCA12 DEGs with expression alteration in other cerebellar ataxias models

In SCA12 fly model, increase activity of pro-apoptotic genes and severe mitochondrial fragmentation has been reported (Wang et al., 2011). We observed differential expression of group of genes related to differential activities of mitochondrial functions. These were (1) dysregulation of *NDUFV2, NDUFS5* and *ATP5C1* genes related to mitochondrial electron transport chain and ATP synthesis (2) upregulation of apoptotic genes *DFFA, PRKAR1B CASP7* (3) upregulation of antioxidant genes such as *CYBA* and *SEPP1* among DEGs. The expression of *SOD2* was observed to be induced (2 fold) but was not within statistical q value <0.05 limit (4) Upregulation of *PPARGC1A* (PGC1α), a crucial component of mitochondrial bioenergetics was also observed.

Calcium signaling has been implicated in pathophysiology of spinocerebellar ataxias (Guergueltcheva et al., 2012; Kasumu and Bezprozvanny, 2012). We observed upregulation of *PRKCB, PLCB1, DRD1, GRM1, ADCY9* and downregulation of *PRKCA* and *EGFR* in diseased stage. Similarly glutamate signaling has been observed to be disrupted in various cerebellar ataxia model system and our results also revealed upregulation of *GRID1* and downregulation of *GRIK5, GRIK2 and GRIA3* genes. Down regulation of an insulin-like growth-factor binding protein (IGFBP5) has been reported in SCA1 and SCA7 transgenic mice (Gatchel et al., 2008). We observed upregulated *IGFBP5* in SCA12 neurons.

Another gene ontology open source tools Enrichr (Kuleshov et al., 2016) also revealed enrichment of GO biological process related to immune responses (Supplementary file 1: Table 9). Altogether, our findings suggest that iPSC derived SCA12 neurons display signatures of immune response and neurodegeneration.

### Elevated expression of CAG repeat containing isoforms discovered in RNA sequencing analysis

*PPP2R2B* transcribes into several neuron-specific splice variants which differ in the 5’ UTR region. Updated NCBI Genome Reference Consortium Human Build 38 (GRCh38), predicts 10 splice variants (V) of *PPP2R2B* which can be divisible into i) CAG repeat containing coding transcripts V3 and V10, ii) CAG repeat containing non-coding transcripts V11 and V12 iii) Transcripts without CAG repeat V2, V4, V5, V6, V8 and V9 [http://www.ncbi.nlm.nih.gov/]. From RNA-Seq data we assessed expression pattern of different splice isoforms of *PPP2R2B*, i.e. transcripts containing CAG repeats (CAG repeat expansion mutation is the causal event) and two of the non-CAG containing transcripts (V2 and V4) due to their earlier shown role in mitochondrial mediated pathology in SCA12 fly model (Wang et al., 2011). From RNA-Seq analysis, we observed a differential expression of *PPP2R2B* splice isoforms across three lineage of SCA12 patients and control cells (Fig. 6A). V3 (CAG coding transcript) was observed to be the most abundant transcript across all the lineages of controls and iPSC and neuronal lineages of disease lines. In control-iPSC, V3 was the only transcript with FPKM >1, in control-NSC expression of other CAG containing transcript (RNA26203 and V10) were observed next to V3, whereas, in control-neurons expression of V2 was second highest after V3. In disease-iPSCs an increase in expression of RNA26203 was noted (>1 FPKM) (Supplementary file 1: Table. 10). In disease-neurons the abundant transcript observed was RNA26203 followed by V2 and V3. In disease-neurons, a slight upregulation of RNA26203 (log2FC 0.78, p-value 0.08) transcript was observed whereas V3 transcripts (log2FC -1.78, p-value 5x10^-^5^^) and V2 (log2FC -0.76, p-value 0.0077) were found downregulated. The expression pattern of *PPP2R2B* isoforms was validated by qPCR and in comparison to control cell lineages, disease lines shown downregulation of V2, V3, V4, V10 and V11, whereas RNA26203/V12 transcript was observed to be upregulated in iPSCs (log2FC 3.5), NSCs (log2FC 4.7) and abundant expression (log2FC 6.07) in disease neurons (Fig. 6B). As RNA-Seq data have not shown any upregulation of RNA26203 in NSCs which was observed through qPCR in disease-NSCs, which prompted us to speculate whether primers (for qPCR) designed for V12 transcript (primers flanked the CAG region) captured any novel isoform, as the expression of V12 was also observed to be highly upregulated in disease neurons. qPCR product, on 2% agarose gel, showed the presence of expanded allelic copy (RNA) along with normal allele (data not shown).

**FIGURE 6:**
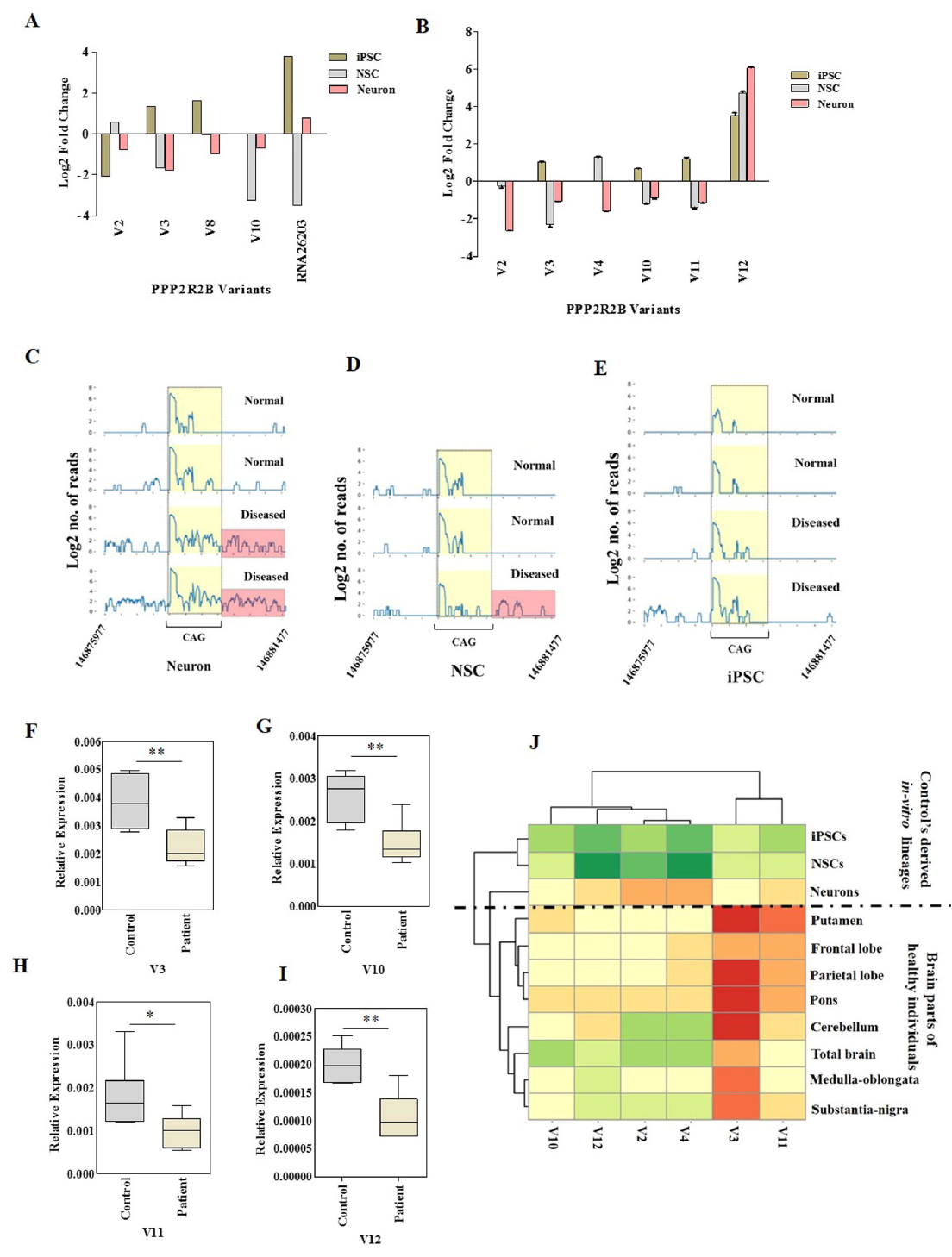
Differential expression of *PPP2R2B* splice variants. **(A)** Quantitative expression (Log2 fold change) of *PPP2R2B* isoforms obtained through RNA sequencing in patient derived neurons, NSCs and iPSCs relative to control (**B)** qPCR based validation of different splice isoforms of *PPP2R2B*. GAPDH was used as internal control for normalization. Relative gene expression was calculated using 2 ^-Δ ΔCt^method and results were plotted as log2 fold change in patient derived lines compared to controls. **(C-E)** RNA sequencing read distribution around CAG repeat region across different cell lineages of patients and controls showing extra reads mapping upstream to the CAG track in patient lines. **(F-I)** qPCR analysis of gene expression for *PPP2R2B* variants V3, V10, V11and V12 in peripheral blood samples of SCA12 patients compared to controls (n=5). GAPDH was used as internal control for normalization and Relative expression was determined through 2 ^-Δ Ct^method. Statistical significance was determined using unpaired t-test (J) Heat map depicts distribution of relative expression (ΔCt) values among several brain regions (control individuals) and in-vitro cell lineages derived from healthy individual. The colour gradient from green to red represents magnitude of expression. (Red represents higher expression whereas green represents lower expression of PPP2R2B). (* = p <0.05, ** =p <0.001).

To further investigate regarding the occurrence of any other isoform of *PPP2R2B*, we additionally aligned RNA-Seq reads for each sample (patient and control lines) for the region around the CAG repeat (chr5:146,877,977-146,879,594) and in the regions near *PPP2R2B* promoter. In this region, we plotted the read density and found that control cell lines from each stage i.e. iPSC, NSC and Neuron contained only a small number of reads upstream of CAG repeat. However, among both the replicates of neuron cells from the disease condition, a uniformly distributed read density could be seen upstream of CAG repeat. This pattern of sequencing read distribution was also observed in NSC stage but not in iPSC. Occurrence of these additional sequencing reads upstream region suggests a possible existence of some novel isoform with respect to *PPP2R2B* in the disease condition (Fig. 6C-E). Further to explore the expression pattern of *PPP2R2B* in SCA12-neurons, we also quantified *PPP2R2B* transcriptional changes in peripheral blood of patients and controls (n=5 each). Compared to controls, all repeat containing isoform had lower expression in blood similar to iPSC derived neurons. Among the various isoforms studied, the expression of isoforms V2 and V4 were not detectable in the blood samples of both controls and patients (Fig. 6 F-I) and no heightened expression of V12 was observed.

### Region specific expression of *PPP2R2B* in healthy brain

Initially we compared *PPP2R2B* expression of the cell lines generated and healthy control brain tissues. The expression of all the isoforms of *PPP2R2B*, was found in healthy control derived neurons similar to healthy brain tissues. To study the regional expression of *PPP2R2B* in brain, we also assessed the expression pattern of *PPP2R2B* splice isoforms in different brain regions of healthy individuals’ viz. cerebellum, frontal lobe, parietal lobe, medulla oblongata, pons, putamen, substantia nigra (Fig. 6J). We observed that *PPP2R2B* isoforms were variably expressed across the brain regions and V3 being the highest in expressed in all the regions. The expressions of non-coding CAG containing transcripts was higher in putamen for V11 and cerebellum and parietal lobe for V12.

### Genome-wide transcriptional profiling of peripheral blood of SCA12 patients

Genome wide gene expression analysis of blood transcriptomics identified 11 DEGs (3 upregulated and 8 down regulated) (Supplementary file 1: Table 11). *CCL23* and *LGALS2* were upregulated and among the downregulated set of genes, *CYP261*, *GPR97*, *HEY1*, *LIMK2*, *PI3*, *SLP1* were the important candidates with a possible role in neurodegeneration.

### Profiling of inflammatory signature in peripheral blood of SCA12 patients

Further to explore interferon signature in peripheral blood of SCA12 patients, we quantified the expression of several genes associated with inflammatory response in peripheral blood of representative SCA12 and controls (n=5). We found significant up regulation of *IFI6* in patients blood compared to controls (Fig. 7A-F).

**FIGURE 7:**
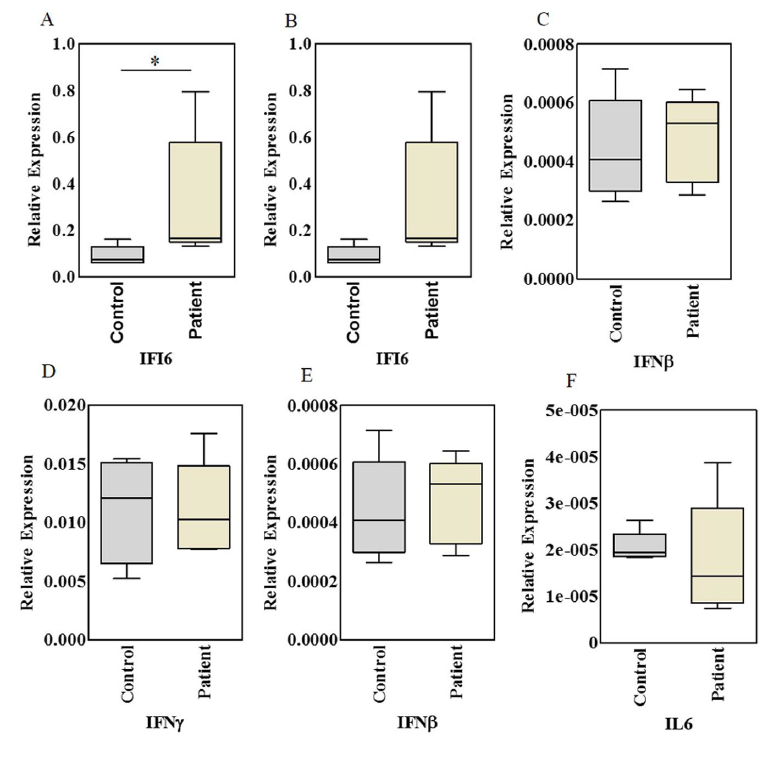
qPCR validation of key interferon siganling pathway genes in SCA12 patients. Relative expression (RE) of selective genes *IFI6, IFI16, IFN*β*, IFN*γ*, Il*β*1, IL6* involved in interferon signaling pathway in peripheral blood samples of SCA12 patients compared to controls (n=5). GAPDH was used as internal control. RE was determined using 2 ^-ΔCt^method. Statistical significance was determined using Mann–Whitney rank sum test. (^*^ =p <0.05)

## Discussion

### Modelling SCA12 using patient-specific iPSCs

Radiologically and neuropathologically, SCA12 brain shows pattern of degeneration restricted to cerebral and cerebellar cortex (Holmes et al., 1999; O’Hearn et al., 2015; Srivastava et al., 2017). An iPSC-derived neuronal lineage might serve as an alternate surrogate experiment material to study the molecular events in SCA12. In our study we were able to generate iPSCs of SCA12 patients noninvasively from LCLs to further differentiate into neuronal lineage. The generated neurons displayed characteristics of a pan-neuronal lineage. We did not expect morphological differences in these SCA12 neurons at early stages as like other neurodegenerative disease SCA12 produces a late onset phenotype and even neuropathological studies have not shown any morphological differences in the disease affected brain areas except for neuronal loss and gliotic changes (O’Hearn et al., 2015).

We however used these derived neurons at early stage for an in-depth characterization at the transcripts level in disease condition of SCA12 and studied the whole transcriptomics and transcription of *PPP2R2B* alternate splice isoforms.

### Signatures of inflammation in both SCA12 derived neuronal lineages and peripheral blood

In our SCA12-neuronal cells, of interests, we observed an induced response of type-I interferon signaling pathway genes (*IFI27*, *IFI6*, *OAS1*, *OAS3*, *MX1* and others). Interferons are a large group of structurally related cytokines play protective defenses against viral infection (Sadler and Williams, 2008). Apart from interference with viral proliferation in a typical scenario induction of interferons can also been observed in the absence of pathogens. IFN activation also play as inflammatory mediators in neurodegeneration in Alzheimer’s disease, Huntington disease and Friedreich Ataxia (FRDA)(Taylor et al., 2014; Miller et al., 2016; Sanchez et al., 2016). Interferon inducible inflammatory responses is frequently linked to interferon-induced apoptosis and DNA damage pathway (Brzostek-Racine et al., 2011). In the present study upregulation of genes involved in DNA damage (*PARP9* and *DTXL3*) and interferon-induced apoptosis (OAS1 and OAS3) were observed and it is consistent with findings reported in FRDA and other neurodegenerative diseases (Nakad and Schumacher, 2016; Sanchez et al., 2016). Stimulation of several interferon induced genes (IFN) such as *OAS1and 3*, *ISG15, MX1* activate latent RNase L, which degrades viral and endogenous RNA and induces transcription of genes related to interferon signaling (Lin et al., 2009; Schneider et al., 2014).

Another gene related to IFN response, *DDX58* and *IFIH1* were found upregulated and of which *DDX58* encode an RIG-I-like receptor dsRNA that recognized viral double stranded RNA (Yoneyama et al., 2004). It initiates immune response through mitochondrial antiviral signaling and shown to be associated with neuroinflammation (de Rivero Vaccari et al., 2014; MacNair et al., 2016). Upregulation of *DDX58* and *IFIH1* in SCA12 suggest that mechanism of neurodegeneration could be a consequence of inflammation that is activated through the RIG1 signaling in response to over expression of RNA.

From peripheral blood transcriptomics data we were able to see an alteration in expression of genes related to immune response. The significantly expressed genes in blood transcriptome of SCA12 belonged to innate immunity. Among these, *CCL23* is a member of the CC chemokine subfamily and *GPR97* belongs to adhesion G-protein coupled receptor family reported as biomarker in inflammatory diseases (Pawlak et al., 2014). Similarly, *LIMK2* is a member of PDZ/LIM family which has role in physiological functioning of nervous system pro-survival induced by DNA damage pathway and genotoxic stress (Cuberos et al., 2015). Among other DEGs, *PI3* and *SLP1* are related to innate immune response and their synergistic functions play an antimicrobial role (Sallenave, 2010). Another candidate of interest, *IFI6 b*elong to family of interferon inducible factor and play role in interferon signaling pathway. Its encoded protein play role in antiviral activity and regulation of apoptosis (Meyer et al., 2015; Qi et al., 2015). Upregulation of *IFI6* in peripheral blood similar to RNA-Seq findings further supports the involvement of immune mechanism in pathogenesis of SCA12. Inflammation linked to antiviral immune response through RNA triggers is implicated in pathogenesis of diseases (Olejniczak et al., 2015). The secondary structure of expanded RNA and mutant transcripts, products of bidirectional transcription, alternatively spliced transcripts, and RNAs released from necrotic cells are recognized by pathogen-associated molecular patterns receptors and cellular sensors (PKR, OAS, RIG-I, MDA5 and Toll-like receptors) (Tian et al., 2000; Bañez-Coronel et al., 2012; Nalavade et al., 2013; Richards et al., 2013). This may activate several interferon induced genes OAS family resulting into transcription of inflammatory genes. Though, at this point we are not able to show the formation of these RNA secondary structures in SCA12-neurons or other lineages but RNA mediated inflammatory mechanism is speculative in SCA12 (Nalavade et al., 2013; Olejniczak et al., 2015). Our whole transcriptomic studies on SCA12-neuron suggested an induced response to IFN signaling and possible indicating and RNA driven mechanism of these inflammatory signals. The observation of a highly upregulated non-coding CAG containing V12/RNA26203 or yet unidentified novel isoform of *PPP2R2B* in our study supports this possibility, while all other transcript isoforms of *PPP2R2B* were found downregulated.

### Recapitulation of gene expression signatures of other neurodegenerative diseases in SCA12 iPSC-derived neurons

In addition to inflammatory response, our study has observed dysregulation of key pathways implicated in neurodegeneration and other related cellular processes. We observed aberrant expression of genes related to mitochondrial bioenergetics, NADPH (Nicotinamide adenine dinucleotide phosphate) and cytochrome oxidase complex components, apoptosis and antioxidant. This corroborates with the observation in SCA12 fly model (Bβ2 twins/tws overexpression model) wherein mitochondrial fragmentation with cristae disruption were observed in photoreceptor neurons of SCA12 fly, accompanied with increase in reactive oxygen species (ROS) production, cytochrome c, and caspase 3 activity leading to apoptosis and decreased life span of fly (Dagda et al., 2008; Wang et al., 2011). We also observed an upregulation *PPARGC1A* (PGC1α), a crucial component of mitochondrial bioenergetics similar to as reported in Parkinson disease (Zheng et al., 2010). Notably, the calcium and glutamate signaling has been observed to be disrupted in various cerebellar ataxia model system. In the present study we also identified several differentially genes belonging to such important pathways. Dysregulation of insulin-like growth-factor pathway has reported in SCA1 and SCA7 (Gatchel et al., 2008) and we observed an upregulation of *IGBP5* in SCA12 neurons. Of interest, among pathways of other neurodegenerative disorders, we observed some overlap of DEGs with genes reported in Alzheimer disease (*BACE2*), and mitochondrial genes *NDUFV2* and *NDUFS5* common to AD, PD and HD pathways (Supplementary file 1: Table 8). This suggests a common link of neurodegeneration through our SCA12-neuronal cells.

In the existing literature it has been reported that *PPP2R2B* is brain specific in expression *PPP2R2B* gene encodes a regulatory subunit of PP2A, known to be involved in several inter-related signaling pathways (Janssens and Goris, 2001; Dagda et al., 2003; Sangodkar et al., 2016). The aberrant expression of *PPP2R2B* may affects PP2A activity functioning which in turn dysregulate various signaling cascade as we have observed an overall downregulation of *PPP2R2B* might result in a loss of function. RNA mediated toxicity has been implicated in many TNR diseases caused by repeat expansion either in the coding or non-coding region (Krzyzosiak et al., 2012). A similar scenario can be anticipated in SCA12 where the CAG repeat expansion in a non-coding (untranslated) region of *PPP2R2B.* In SCA12 it has been shown that the length of the CAG repeats alters the expression of the *PPP2R2B* (Lin et al., 2010; O’Hearn et al., 2015). Overexpression of CAG repeat containing isoform of a non-coding RNA in SCA12 is reminiscent of the expression pattern observed in FXTAS (Hagerman, 2013) and may exert cytotoxic effects through various mechanisms such as RAN translation, antisense and poly-serine mediated pathophysiology (Zu et al., 2011; Nalavade et al., 2013). Elongated CAG repeat motifs in their UTRs form secondary structures which can sequester different important proteins, leading to the formation of RNA foci and subsequently causing cell death. Ubiquitin positive inclusion bodies in the substantia nigra have also been observed in SCA12 (O’Hearn et al., 2015). Hence, there might exist overlaps in the disease mechanism between SCA12 and FXTAS.

## Conclusions

In SCA12-neuronal cells, we have observed a heightened expression of CAG expanded RNA transcript in neuronal lineages. The identified signatures of inflammation in the peripheral blood tissue and patient iPSC-derived neurons open the possibilities of monitoring the progression and prognosis of disease. This may be helpful to identify anti-inflammatory molecules that can be effective as a therapeutic approach or for slowing the progression of disease. To our knowledge, this is the first study to address SCA12 pathogenesis using patient derived iPSC/neuronal cells and. and generates transcriptomic data resources for future studies.

## Abbreviations

ALS: Amyotrophic lateral sclerosis
FC: Fold Change
FDR: False Discovery Rate, FPKM, Fragments Per Kilobase of transcript per Million mapped reads
FRDA: Friedreich Ataxia
FXTAS: Fragile X-associated tremor/ataxia syndrome
HD: Huntington’s disease
iPSCs: Induced Pluripotent Stem Cells
LCLs: Lymphoblastoid Cell lines
non TNR: non trinucleotide repeat
NSCs: Neural stem cells
PD: Parkinson’s disease
qPCR: Quantititave PCR
RNA-Seq: RNA sequencing, SCA12, Spinocerebellar ataxia type 12
TNR: Trinucleotide repeat

## Conflict of interests

The authors declare no competing financial interests.

### Acknowledgements

This study was supported by Council of Scientific & Industrial Research (CSIR) funded project GENCODE (BSC0123). For OM, funding from Shanta Wadhwani Centre for Cardiac and Neural Research (SWCCNR) and InStem is acknowledged. Funding from UGC for AH is duly acknowledged. The authors thank all the subjects who had participated in this study. We acknowledge Varun Suroliya, Renu Kumari for assistance in RNA-Seq experiments and Rajesh Pandey for his assistance in microarray experiment.

## Authors’ contributions

DK, MF, conceived and designed the study. DK performed experiments. PD, RK, MF analysed transcriptomic data. AH, OM supported experiments. AKS provided support in recruitment of subjects, AKS and MF helped in clinical evaluation of patients. DK, MM, MF prepared the manuscript. All authors read and approved the final version of the manuscript for submission.

